# Improving hifiasm haplotypes for autopolyploid genome assemblies using constraint programming

**DOI:** 10.1101/2025.04.01.646355

**Authors:** Christophe Klopp, Valentin Durante, Thomas Schiex, Simon de Givry

**Affiliations:** Université Fédérale de Toulouse, INRAE, Sigenae, MIAT, BioinfOmics, Chemin de borde rouge, 31326, Castanet-Tolosan, France; Université Fédérale de Toulouse, INRAE, SaAB, MIAT, Chemin de borde rouge, 31326, Castanet-Tolosan, France; 5 Degrés Sud, 8 Grande rue de nazareth, 31000, Toulouse, France

**Keywords:** autopolyploid, haplotypic genome assembly, constraint programming

## Abstract

Actual versions of hifiasm often produce unbalanced haplotype contigs for autopolyploid genome assemblies. Adding a contraint on the distribution of proteins among haplotypes improves haplotype sizes and busco scores balances. Four species were used to benchmark this approach against hifiasm raw results. The script can be downloaded from https://github.com/chklopp/Separate_haplotypes_with_toulbar2.

## Introduction

Today high quality long read sequencing technologies produce data enabling to easily assemble large genomes. Increased read length and lowered error rates make it possible to assemble large repeats harboring only few variations between copies. These improvements push toward generating more complete assemblies such as haplotypic or T2T and also to assemble more difficult genomes such as polyploids and particularly autopolyplöids. Portions of the genome can be specific to one haplotype, shared by several or shared by all haplotypes. This renders the assembly more difficult as the assembly graph gets more complicated. hifiasm [3] has a `--n-hap` parameter enabling to output more than two haplotypes. This parameter can only be used if Hi-C reads are provided to the software package. In this case it will output as many assembly files as the number of defined haplotypes. Unfortunately the outputs are often not balanced and the BUSCO [12] scores show wrongly placed contigs meaning that one or several haplotypes have duplications and others have missing BUSCO proteins. Before separating contigs in haplotypes, it is important to check that the merged set of contigs harbors the kmers found in the reads with the correct number of copies. This can be done using kat [8] spectra-cn plot to visually inspect how number of copies are distributed among the expected kmer counts.

Constraint programming is an artificial intelligence technique enabling to describe a problem with a set of finite-domain variables and a set of cost functions [Handbook of constraint programming [1]]. Each constraint involves a subset of variables called its scope. A constraint describes a list of allowed assignments for the variables in its scope. A cost function is a function which associates a cost to each possible assignment of its variables. The goal is to find a complete assignment which satisfies all the constraints and minimizes the total cost of all the cost functions.

Hereafter we present `shbt2.py` a python script which uses the haplotypic contig sets provided by hifiasm and constraints generated from protein alignments on these contigs in order to generate novel haplotypes which are more balanced in term of total size and BUSCO scores than the initial hifiasm results. We will describe processing steps, constraints and results on four different public data sets to validate the haplotypic metric improvements.

### shbt2.py script

The `shbt2.py` script chains all the processing steps described in the next paragraph and produces haplotypic assembled multi-fasta files.

#### processing steps

The script first verifies that the needed software packages hence xz, miniprot [7] and toulbar2 [5, 9] are available in the environment. If this is the case it aligns the proteins to the contigs using miniprot allowing a maximum of alignments per protein corresponding to the species ploidy. From the alignment file it generates a contig to protein link file which is used to build the first constraint set. If a contig to haplotype link file is given then a second set of constraints is added. The constraint file is then processed by toulbar2 to produce a solution file containing a list of contigs per haplotype. Last, the solution file is processed to produce haplotypic multi-fasta files.

#### Constraints description

The script can work with one or two constraint sets. The first mandatory cot function set is built from the protein links of the contigs. The cost of putting two contigs in the same haplotype is the square of the number of shared proteins. The cost of the second cost function is nil if the contig is placed in its initial haplotype from hifiasm and the number of proteins it contains if it is placed in another haplotype.

#### script options

The script has three mandatory options which are the contig multi-fasta file (`--assembly`), the protein multi-fasta file (`--proteins`) and the species ploidy (`--ploidy`). The second set of cost functions is added by the `--groups` parameter which corresponds to a tabulated file in which the first column contains the contig names and the second the group indices starting from zero. The `--optime` parameter enables to give the processing time limit for toulbar2 to produce the solution for this NP-hard problem.

### Benchmark methods and results

#### public data sets

Four public data sets were used for benchmarking. They correspond to four different species: *Actinidia arguta, Fragaria x ananassa, Medicago sativa* and *Solanum tuberosum*. The expected haplotype assembly size, ploidy, Hifi and Hi-C reads sets are presented in Table 1. All but the strawberry have been assembled and presented in the following publications [6, 2, 11]. The strawberry data without assembly has been published in [4].

**Table 1.** public data sets.

| Species | S (Mb) | Pl | Hifi acc. | Hi-C acc. |
| --- | --- | --- | --- | --- |
| *A. arguta* | 700 | 4 | SRR24888632 | SRR24888631 |
| *Fxa F. brilliance* | 240 | 8 | SRR21850459 | SRR21850458 |
| *M. sativa* | 780 | 4 | SRR11285798 | SRR13333748 |
| *S. tuberosum* | 850 | 4 | SRR1681122[6-8] | SRR15560098 |

The data sets have been downloaded from the NCBI sra website. "S" corresponds to the size of each haplotype. "Pl" corresponds to genome ploidy. "Hifi acc." and "Hi-C acc." correspond to all the accessions used for the benchmark.

#### Processing steps

Hifi and hi-C reads were assembled with hifiasm version 0.24.0 using the `--n-hap, --hom-cov, --hg-size, --h1` and `--h2` parameters. The hapotype assemblies metrics were calculated with `assemblathon stats.pl` and checked with BUSCO [12] 5.4.7 with the embryophyta odb10 reference database. All the haplotyped assemblies were merged in a single assembly file which was then checked for kmer completeness with merqury [10] version 1.3. The total assembly length, contig N50 and merqury completeness are presented in Table 2. For each species a haploid protein set of a phylogenetically closely related species (*Actinidia eriantha* for *Actinidia arguta, Fragaria vesca* for *Fragaria x ananassa, Medicago truncatula* for *Medicago sativa* and *Solanum lycopersicum* for *Solanum tuberosum*) was downloaded from the NCBI and `shbt2.py` was ran giving the merged haplotypes produced by hifiasm and the protein file. Hifiasm and shtb2 haplotypes were compared and results are presented in next paragraphs.

**Table 2.** hifiasm merged assembly metrics.

| Species | Total length | contig N50 | Completeness |
| --- | --- | --- | --- |
| *A. arguta* | 2.9 | 6.25 | 98.75 |
| *Fxa F. brilliance* | 2.1 | 4.43 | 98.42 |
| *M. sativa* | 3.3 | 1.39 | 96.82 |
| *S. tuberosum* | 3.2 | 6.95 | 98.76 |

"Tot length" corresponds to the total size of all haplotypes merged. "contig N50" and merquy "Completeness" have been calculated for all haplotypes merged.

#### haplotype size comparison

Figure 1 shows the difference in total contig length of the haplotypes generated by hifiasm and the ones produced by `shtb2.py`. These are much more balanced particularly for *Fragaria x ananassa*.

**Fig. 1.**
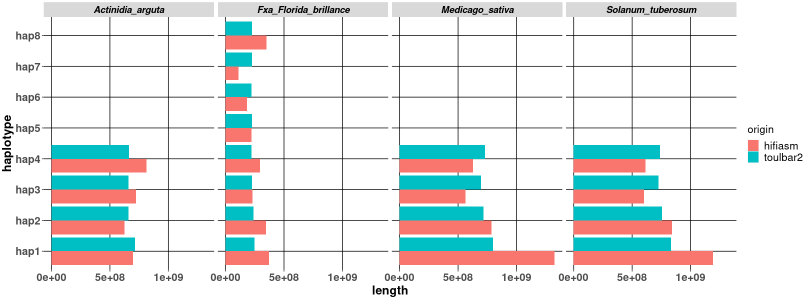
For each species and each haplotype this figure presents the hifiasm, in orange, and toulbar2 in blue, total haplotype contig length. Toulbar2 produces better balances haplotypes.

#### BUSCO score comparison

Figure 2 shows the BUSCO metrics of the haplotypes generated by hifiasm and the ones produced by `shtb2.py`. These are much more balanced which is expected for autopolyploids in which each haplotype can be equally transmitted to the offspring.

**Fig. 2.**
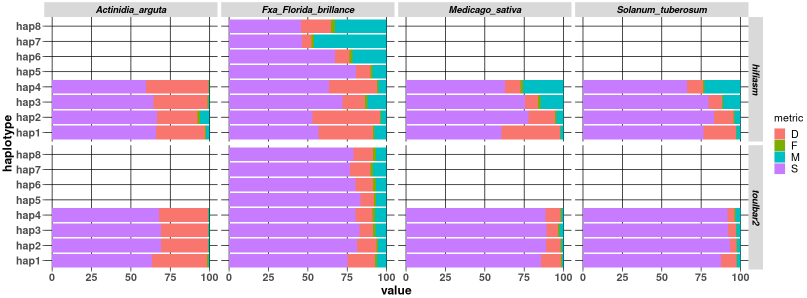
For each species and each haplotype this figure presents the hifiasm at the top, and toulbar2 at the bottom BUSCO metrics. S: single, D: duplicate, F: fragmented or partial protein, M: missing on not found protein

## Conclusion

Being able to build all haplotypes of an autopolyploid species is very important to access variability. Hifiasm is a valuable help in autopolyploid genome assembly but its outputs do not comply with the expected haplotype balance. Constraint programming enables to merge initial assembly results with protein content constraints to improve haplotypes. Other functions, related for example to Hi-C links, could be added to the model to further improve the final result.

## Author contributions statement

T.S., V.D., S.d.G and C.K. conceived the script, C.K. conducted the benchmark, T.S. S.d.G and C.K. analysed the results. V.D. T.S. S.d.G and C.K. wrote and reviewed the manuscript.

## Acknowledgment

We are grateful to the genotoul bioinformatics platform Toulouse Occitanie (https://doi.org/10.15454/1.5572369328961167E12) for providing help and/or computing and/or storage resources. We thank Adela Poublan-Couzardot and Christine Gaspin for the fruitfull discussion we shared on the subject.

